# Pragmatic Language Processing in the Adolescent Brain

**DOI:** 10.1101/871343

**Authors:** Salomi S. Asaridou, Ö. Ece Demir-Lira, Julia Uddén, Susan Goldin-Meadow, Steven L. Small

## Abstract

Adolescence is a developmental period in which social interactions become increasingly important. Successful social interactions rely heavily on pragmatic competence, the appropriate use of language in different social contexts, a skill that is still developing in adolescence. In the present study, we used fMRI to characterize the brain networks underlying pragmatic language processing in typically developing adolescents. We used an indirect speech paradigm whereby participants were presented with question/answer dialogues in which the meaning of the answer had to be inferred from the context, in this case the preceding question. Participants were presented with three types of answers: (1) direct replies, i.e., simple answers to open-ended questions, (2) indirect informative replies, i.e., answers in which the speaker’s intention was to add more information to a yes/no question, and (3) indirect affective replies, i.e., answers in which the speaker’s intention was to express polite refusals, negative opinions or to save face in response to an emotionally charged question. We found that indirect affective replies elicited the strongest response in brain areas associated with language comprehension (superior temporal gyri), theory of mind (medial prefrontal cortex, temporo-parietal junction, and precuneus), and attention/working memory (inferior frontal gyri). The increased activation to indirect affective as opposed to indirect informative and direct replies potentially reflects the high salience of opinions and perspectives of others in adolescence. Our results add to previous findings on socio-cognitive processing in adolescents and extend them to pragmatic language comprehension.

## INTRODUCTION

Adolescence is a period during which social interactions, especially with peers, become increasingly important [17, 20, 23, 33]. Pragmatic competence, defined as the appropriate use of language in social situations, is crucial for successful social interactions. This ability is manifested in conversational turn taking, the ability to maintain topic relevance, express requests or declinations, understand utterances based on context, and adjust language based on the communicative situation (for instance, knowing when and how to use slang expressions) [50]. It is therefore not surprising that pragmatic competence strongly influences peer acceptance in adolescence. Ten-year-old children who showed pragmatic competence during a conversation were judged as more popular, attractive, and academically successful by their peers [44]; shy preschool children were judged as more peer-likeable by their teachers when they showed good pragmatic skills [14]; and unpopular preadolescent children improved their peer acceptance and self-perceived social efficacy after receiving conversational skills training [7].

Pragmatic competence starts developing very early [4, 12] and is likely to continue throughout adolescence, although this has rarely been studied empirically (see however [19, 50]). Aside from core language abilities (such as vocabulary and grammar), theory of mind (ToM), i.e. the ability to attribute thoughts, perspectives, opinions, and emotions to others, contributes to the emergence of pragmatic competence during development [4, 37, 64]. ToM is still developing in adolescence; adolescents make more errors and use perspective taking less frequently than adults when performing tasks that require representing other people’s mental states [19, 29, 57, 59].

Adolescence is not only a period of rapid behavioral change in social skills but is also characterized by significant neuroanatomical changes. These changes include accelerated decreases in cortical thickness in frontal, temporal and parietal areas [58]. Changes in these particular areas are noteworthy since they are thought to play a role in ToM development and might be contributing to the development of pragmatic skills.

Despite its behavioral significance and possible links to neurocognitive changes, little is known about the neural underpinnings of pragmatic language understanding in adolescents. Here we aimed to identify the brain networks recruited by adolescent listeners during discourse comprehension. We used dialogues consisting of questions and answers between interlocutors. The answers could be unambiguous, literal replies to the question (henceforth direct replies) or non-literal replies (indirect replies). The dialogues were constructed in such a way that the directness of the replies was manipulated solely by changing the queries, i.e., direct and indirect replies shared identical text. An example of a *direct* reply dialogue was: Q: “Where should we go for a nice family vacation?” A: “Disneyland is a great place for little kids”. For indirect replies, the listeners would have to infer the intended meaning from the context. There were two types of indirect replies, each with its own specific type of ambiguity: *informative* replies provided extra information to what would otherwise be a yes/no reply (Q: “Do you think the children will have fun on the trip?” A: “Disneyland is a great place for little kids”), while *affective* replies used ambiguity to politely convey negative opinions, refuse requests or save face (Q: “Wouldn’t it be great to go to Disneyland for our honeymoon?” A: “Disneyland is a great place for little kids”; see also Fig. 1). The listener had to infer the speaker’s intention by taking into account the content of the preceding question, as well as the speaker’s voice, prosody, and/or the relationship between interlocutors. The process of interpreting meaning via context-dependent inference offers a measure of pragmatic abilities that is distinct from core language abilities [67].

**FIG. 1.**
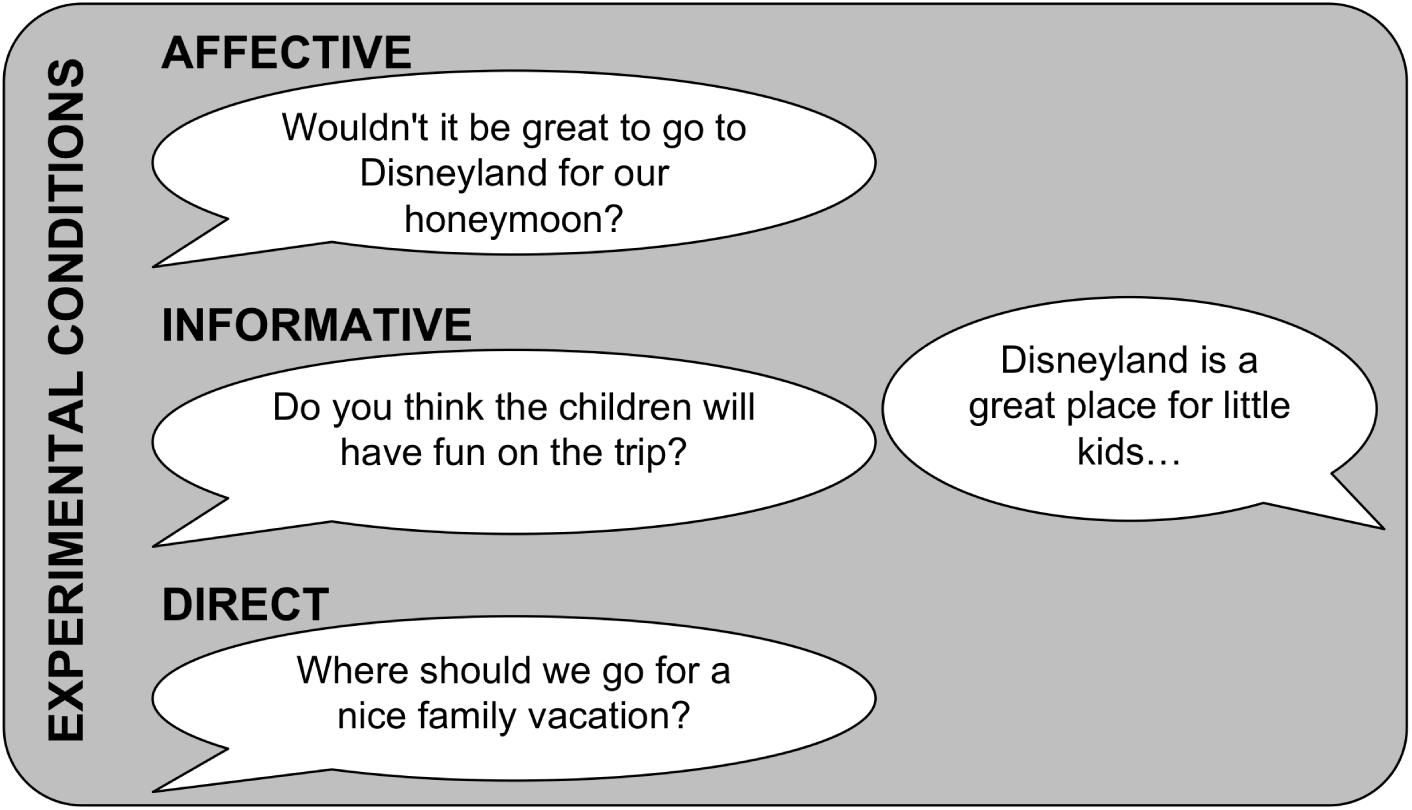
Example question – answer dialogues in each experimental condition.

Our first hypothesis was that listening to the indirect replies (whether informative or affective) would require listeners to consider the literal meaning of the reply and make inferences about the intended meaning [26]. Indirect replies would consequently engage a more extended brain network than direct replies, as has been previously found in adult listeners [6, 55, 60, 66]. Specifically, we expected higher activity in certain language processing areas (STG/STS in both hemispheres), and potentially in brain areas associated with attention, including the superior and inferior parietal lobes, and working memory, including the ventrolateral PFC. Our second hypothesis was that interpreting the meaning of the indirect affective replies would require ToM skills, and would thus lead to higher activity in the medial prefrontal cortex (mPFC) and the posterior superior temporal sulci/angular gyri (temporoparietal junction/TPJ). We reasoned that indirect replies would activate regions known to be active in adolescents during social-cognitive tasks, as shown in cross-sectional comparisons [11, 13, 34, 53]. Given adolescents’ heightened sensitivity to social acceptance, we anticipated that affective replies would elicit more salient and pronounced activation compared to informative and direct replies in regions of the ToM network. Lastly, although processing affective replies involves inferring speaker’s intention and revising the reply meaning, we did not expect listeners to experience the emotions of the speaker or receiver of the reply. Thus, no increased activity was expected in limbic network areas (amygdala and/or anterior insulae), especially as participants were overhearers and not addressees of the dialogues.

## METHODS

### Participants

Twenty-five typically developing adolescent children (age 14-17 years, mean age ± SD: 15*y*; 9*mo* ± 8.2*m*; 3 left-handed; 10 female) took part in the study. Participants were part of a larger longitudinal cohort selected to be demographically representative of the greater Chicago area. Parents gave written informed consent following the guidelines of the Institutional Review Boards for the Division of Biological Sciences at The University of Chicago, and the Office of Research at the University of California, Irvine, which approved the study. Children gave verbal assent. All participants were monolingual native speakers of American English, reported normal hearing and normal or corrected-to-normal vision, and had no history of neurological or developmental disorders.

### Stimuli

Our stimuli consisted of 324 spoken question-answer dialogues - 108 answers, each following three different types of questions (Fig. 1). A number of dialogues were adapted from Holtgraves [26] and Basnakova et al. [6]. Due to scanning time constraints, the dialogues were not preceded by lead-in context stories (as in Basnakova et al. [6]). Instead, the questions provided the sole conversational context for interpreting the answers and were constructed as follows: questions in the direct condition where all open-ended “wh” questions, questions in the informative condition were closed-ended questions that could be answered with “yes” or “no”, while questions in the affective condition were also closed-ended questions but they differed from the informative ones in that the person asking had something at stake. In both informative and affective conditions, the reply was indirect but the reason for indirectness in the affective condition was to be polite, to avoid disagreeing with, hurting or insulting the interlocutor or to save one’s own face as opposed to just providing more information in the informative condition. We also included 50 filler dialogues which were either short questions answered with a simple yes or no, or two declarative statements. (Stimulus material is available at https://github.com/savvatia/PragmaticLanguageAdolescentBrain/tree/master/stimuli)

Before recording the stimuli, we asked 9 young adults to read the written dialogues and paraphrase the answer; items that were hard to paraphrase or misinterpreted were edited or excluded. Participants performed above chance and correctly interpreted affective answers 89.09% of the time, informative ones 86.07% of the time, and direct ones 85.21%. This is quite remarkable, especially given that it is harder to get the gist of a dialogue in written form without intonation, pitch, or speech rate cues. When confused, these young adults misinterpreted the affective answers most frequently as informative, the informative as affective, and the direct as affective or informative.

Six young adults (3 male), native speakers of American English, were recorded while reading the dialogues aloud. Speakers were informed about the different conditions and asked to read aloud the dialogues as naturally as possible keeping in mind the specific conversational context. Each dialogue was recorded twice and the best version was chosen for the experiment. All dialogues were recorded in a soundproof booth using a sampling rate of 44.1 kHz with GarageBand digital audio workstation and their sound amplitude was normalized. Dialogues in each condition did not differ in terms of number of words, word frequency, or concreteness. They did differ in speech rate and intonation, with affective dialogues being uttered at a slower speech rate and more varied intonation. As a result, affective replies were longer in duration (mean=2.90s, SD=0.94) than the informative (mean=2.41s, SD=0.69) and the direct (mean=2.33s, SD=0.65) replies. The pause between question and answer also differed per condition with a longer pause in the indirect (affective and informative) conditions than the direct. These differences in speech rate, intonation, and pause duration were retained because they provided listeners with important cues for understanding the communicative context in which the dialogues were taking place. Normalizing these parameters or using the same recording for the reply across conditions would have introduced unnaturalness in the stimuli which we wanted to avoid. Lastly, there was significantly greater content word overlap between questions and answers (percent of content words in critical utterance that subject has already heard in the question) in the direct condition than in the other two conditions.

### fMRI Task

Participants were told that they would hear everyday dialogues between friends, family members, and colleagues and were asked to listen carefully and try to understand the meaning conveyed by the speakers. We used an event-related design with trial order and timing optimized using optseq2 (https://surfer.nmr.mgh.harvard.edu/optseq). Stimuli were distributed across three lists in which each reply was presented in a different condition so that no participant would hear the same reply in two different conditions. Each list consisted of 36 dialogues per condition, 30 filler items and 13 catch-trial questions. The presentation order was pseudorandomized such that participants heard no more than 2 dialogues from the same condition consecutively. Each trial would consist of a dialogue followed by a jittered inter-trial interval (fixation cross). Catch trials appeared only after filler items, were accompanied by a ringtone, and required participants to read a simple true or false question such as “The speakers mentioned X” and answer by pressing a button with their right hand. We used Presentation software to present the stimuli and record responses (www.neurobs.com).

### Data Acquisition

MRI data were collected on a 3T Siemens Prisma Scanner with a 32-channel head-coil at Northwestern University’s Center for Translational Imaging in Chicago on a single visit. A T1-weighted structural scan was acquired with a magnetization-prepared rapid gradient echo (MP-RAGE) sequence (TR = 2300 ms, TE = 1.86 ms, flip angle = 7°, Inversion Time = 1180 ms, 208 contiguous sagittal slices, slice thickness = 0.8 mm, voxel size = 0.8 × 0.8 × 0.8 mm3, matrix size = 320 × 320). Task fMRI data were acquired with a T2*-weighted echo planar imaging (EPI) sequence with TR = 2000 ms, TE = 25 ms, flip angle = 80°, and 64 axial slices in interleaved order (slice thickness = 2 mm, voxel size = 2 × 2 × 2 mm3, matrix size = 832 × 768). Head motion was minimized using foam padding around the head, and scanner noise was minimized with earplugs.

### Post-scanning questionnaire

The forced-choice post-scanning recognition questionnaire consisted of 27 trials (9 per fMRI task condition). Participants were presented with two written sentences on the computer screen; one of the sentences had been presented inside the scanner (a reply to a question) while the other was completely new. The new sentences were constructed to resemble the ones in the scanner. Participants were asked to choose the one they recognized from the scanner.

### Empathy Quotient

We used the items from the Empathy Quotient questionnaire for adolescents [2] but asked the children to complete the questions instead of their parents. Each question was presented on a computer screen and consisted of a statement about real life situations and experiences. Participants were asked to reply using a 4-point Likert scale (Strongly agree, Slightly agree, Slightly disagree, Strongly disagree). The responses were scored following [2] whereby higher scores indicate higher social and empathy skills. We hypothesized that the skills measured with this questionnaire will be associated with brain activity during pragmatic language processing, particularly the affective condition.

### Debriefing questions

At the end of their visit, participants were asked to answer four debriefing questions, namely to rate their motivation, how interesting they found the study, and how likely it was that they had fallen asleep on a scale from 1 (not motivated/not interesting/not likely) to 4 (very motivated/very interesting/very likely). The fourth question asked them to report any comments or anything they noticed and would like to share.

### Procedure

Data were collected on a single visit. After providing verbal assent, participants went through MRI safety screening and completed the short task training before going into the scanner. A structural scan was acquired first followed by 10 minutes of resting state fMRI during which participants were instructed to look at a fixation cross. A brief sound calibration was performed before starting the task in order to ensure that the sound was at a comfortable level and that each participant could hear the stimuli clearly above scanner noise. Task fMRI data were acquired over three separate runs lasting around 9-10 minutes each. After the task runs were completed, 5 more minutes of resting state fMRI were acquired. Lastly, diffusion weighted imaging data were acquired. Resting-state and diffusion data will not be discussed here. At the end of the imaging session, participants completed the post-scanning questionnaire, the Empathy Quotient questionnaire, and answered the debriefing questions. The whole visit lasted approximately two and a half hours.

### fMRI Analysis

fMRI data were analyzed using AFNI (Analysis of Functional Neuroimages, Version AFNI 17.3.07; http://afni.nimh.nih.gov/ [15, 16]). Dicom files were converted to AFNI BRIK and HEAD files (using the AFNI program Dimon) and the first 3 TRs from every run were removed (3dTcat). Outlier volumes, defined as the volumes with a 0.1 or larger fraction of voxels whose signal intensity deviated from the median absolute deviation of the voxels in the total volumes across runs, were detected (3dToutcount) and the volume with the fewest outlier voxels was used as the base volume for registration. Next, spikes were removed (3dDespike) and slice-time correction with Fourier interpolation (3dTshift) applied. The T1 structural volume was skull-stripped (3dSkull-Strip) and warped to the obliqueness of the functional images (3dWarp). The functional volumes were warped in standard MNI space using the following steps: 1) the transformation matrix from T1 structural space to the functional base volume was estimated (align epi anat.py) and inverted, 2) the T1 structural image was warped to MNI space (@auto tlrc) using a pediatric MNI template (nihpd pediatric asym 13 18.nii.gz), 3) the co-registration of all the functional volumes to the EPI base volume was estimated using affine transformation with a cubic polynomial interpolation (3dvolreg), and finally 4) the transformation matrices from steps 1, 2, 3 were concatenated and applied in a single step on the functional data (3dAllineate). The functional volumes were then smoothed at Full-Width Half Maximum using a 6mm kernel (3dBlurToFWHM) and scaled such that each voxel’s time series had a mean of 100 (3dcalc). The T1 structural volume was segmented into grey matter, white matter and CSF (3dSeg), eroded ROI masks were created for each class of tissue (3dmask tool), and the average time-course for white matter and CSF was extracted (3dmaskave). Motion outliers (volumes with Euclidean norm of motion derivatives exceeding 0.2 mm) were detected (1d tool.py) and combined with the intensity outliers into a single censor regressor for first level analysis (1deval). Participants were excluded from group analysis if more than 30% of their time points were censored. We modeled the hemodynamic response by convolving the stimulus presentation design matrix with a duration modulated block function (dmBLOCK); the resulting regressors for each condition (direct, informative, affective questions and replies) were used as predictors in a general linear model (GLM). Nuisance regressors, including trials of no interest (instructions, filler trials, catch trials and button presses), six motion parameters, average white matter and CSF signal, were included in the model along with four polynomial terms modeling the baseline (constant) and drifts (linear, quadratic, cubic, quartic). Outliers (motion and intensity) were censored at this step. The temporal signal to noise ratio (tSNR) was estimated for each participant using the standard deviation of the GLM error time series (3dTstat/3dcalc). The beta weights from the GLM for the contrasts between the three conditions of interest ([affective replies ¿ baseline] vs. [direct replies ¿ baseline], [affective replies > baseline] vs. [informative replies > baseline], and [informative replies > baseline] vs. [direct replies > baseline]) for each subject were entered in three separate one sample t-tests (3dttest++), one for each contrast. The resulting maps were thresholded using the 3dttest++ “-Clustsim” option which uses the test’s residuals to simulate null 3D results and define cluster size based on them (voxel-wise p=.001, FWE=.05, two-sided thresholding, nearest neighbor=1 [faces touching]). Four additional t-tests were performed on the [affective replies > baseline] vs. [direct replies > baseline] and [affective replies > baseline] vs. [informative replies > baseline] contrasts in which post scanning dialogue recognition accuracy scores and empathy quotient scores were included as covariates.

## RESULTS

### Behavioral Results

All participants responded above chance to in-scanner catch trial questions (*M* = 97.07% correct, *SD* = 7.72%, *range* = 66.67% **–** 100%) and to the post-scanning recognition questionnaire (*M* = 87.26% correct, *SD* = 9.63%, *range* = 55.56% **–** 100%), and the two measures were highly correlated (*r*(22) = 0.67, *p* = .0003) indicating that both measured attention to the dialogues in the scanner. Affective and non-affective (informative and direct) items were equally well remembered in the post-scanning test [Affective: *M* = 88%, *SD* = 11.53%; Non-affective: *M* = 86.89%, *SD* = 10.25%; *t*(24) = .56, *p* = .58]. The empathy quotient scores (*M* = 38.88, *SD* = 9.47, *range* = 19 –58, maximum attainable = 70) did not differ by gender (*t*(21.95) = .10, *p* = .33) and did not correlate with post-scanning accuracy (*r*(23) = **–**.06, *p* = .79). With respect to the debriefing questionnaire, only one participant commented, indicating that “The dialogues were rather passive aggressive and funny”.

### fMRI Results

We excluded two participants from the group fMRI analysis, one due to excessive motion and one because events were not properly logged due to technical error. The analysis was thus performed on the remaining sample of *N* = 23 participants. Figures 2 and 3 and Table I summarize the regions that were significantly more active for the contrasts between different conditions. For the affective vs. direct comparison, the largest clusters of activity were found in the temporal lobes, encompassing the superior temporal gyri (STG) bilaterally in what would be considered part of the language network [21]. Notably, regions of the ToM network including the dorso-medial PFC (dmPFC) and both angular gyri (AG) were reliably more active for affective replies than direct replies. There was also greater activity for affective vs. direct replies in the right inferior frontal gyrus pars triangularis (IFGtri), a region frequently engaged with increasing working memory demands [41, 49], and the pars orbitalis of the left inferior frontal gyrus (IFGorb), which is engaged with increasing semantic integration demands [3, 68, 70]. Affective replies elicited higher activity in both cerebelli (Crus II) (see Figure 3 B), which is consistent with meta-analytic studies showing cerebellar involvement in language, working memory and social cognition tasks [24, 39]. The comparison between the two types of indirect replies, affective and informative, allowed us to disentangle perspective-taking from the effect of indirectness. Consistent with our hypothesis, there was higher activity in the ToM network, including dmPFC and both angular gyri, similar to the affective vs. direct contrast, but there were additional clusters in the ventro-medial PFC (vmPFC) and the precuneus for affective vs. informative replies. Although we did not expect to find any differences in language related areas for this comparison, both STG were reliably more active in the affective compared to the informative condition. Affective replies elicited higher activity in both cerebelli (Crus II) compared to informative replies, possibly indicating a role in processing social information, as well as in areas involved in semantic comprehension, specifically in both IFGorb and left anterior temporal pole. Lastly, we found higher activity for informative vs. affective replies in the right frontoparietal network (right dorsolateral PFC and superior parietal gyrus) which is associated with multiple cognitive processes [43], such as reasoning [63] and attention [32]. To examine the effect of indirectness, independent of perspective-taking, we compared the informative to the direct replies. The comparison revealed two clusters: one in middle right STG extending to the middle temporal gyrus (MTG) and one more posterior, extending to the planum temporale (see Figure 3 A). We did not find higher engagement of any left fronto-temporal areas or right fronto-parietal areas for the informative vs. direct comparison. We did not find significant effects of the EQ or post scanning recognition accuracy co-variates. However, when either of these was added to the model, the activation cluster in the right AG was no longer present (i.e., significant) for the affective vs. informative and affective vs. direct contrasts. Furthermore, adding EQ as a covariate in the informative vs. affective contrast revealed additional significant clusters in the left middle frontal gyrus (BA 46/10) and left superior parietal gyrus (BA 7).

**TABLE I.** Regions showing reliable differences for the Affective vs. Direct, Affective vs. Informative and Informative vs. Direct group comparisons (thresholded at voxelwise *p* = 0.001, FWE=0.05, NN1). XYZ MNI coordinates refer to center of mass location using the LPI coordinate system.

| Contrast | Anatomical Region | x | y | z | Cluster size | Z | MI |
| --- | --- | --- | --- | --- | --- | --- | --- |
| Affective > Direct |  |  |  |  |  |  |  |
|  | R. Superior Temporal Gyrus | 57.5 | -12.7 | -1.1 | 3866 | 4.13 | 0.65 |
|  | L. Superior Temporal Gyrus | -56.4 | -21.1 | 0.8 | 2992 | 4.05 | 0.54 |
|  | Dorso-medial Prefrontal Cortex | 0.6 | 51.3 | 37.7 | 724 | 3.66 | 0.42 |
|  | L. Cerebellum (Lobule VIIa Crus II) | -24.6 | -81.9 | -36.1 | 388 | 3.90 | 0.29 |
|  | L. Inferior Frontal Gyrus Pars Orbitalis | -48.8 | 27.8 | -10.1 | 231 | 3.72 | 0.48 |
|  | R. Angular Gyrus (TPJ) | 59.3 | -50.1 | 22.6 | 215 | 3.81 | 0.30 |
|  | L. Angular Gyrus (TPJ) | -54.6 | -59.3 | 30.5 | 209 | 3.70 | 0.22 |
|  | R. Cerebellum (Lobule VIIa Crus II) | 24.8 | -80.7 | -35.9 | 188 | 3.66 | 0.26 |
|  | R. Inferior Frontal Gyrus Pars Triangularis | 56.6 | 22.1 | 24.2 | 90 | 3.62 | 0.25 |
|  | R. Inferior Temporal Gyrus | 51.1 | 6.2 | -40.1 | 49 | 3.82 | 0.22 |
| Direct > Affective |  |  |  |  |  |  |  |
|  | Left Middle Frontal Gyrus | -39.4 | 46.4 | 21 | 52 | -3.81 | -0.17 |
| Affective > Informative |  |  |  |  |  |  |  |
|  | R. Superior Temporal Gyrus | 59.3 | -17.6 | -1.8 | 2211 | 3.98 | 0.49 |
|  | L. Superior Temporal Gyrus | -58.4 | -22.1 | 1.5 | 1590 | 3.79 | 0.43 |
|  | Dorso-medial Prefrontal Cortex | 0 | 55 | 29.2 | 920 | 3.82 | 0.41 |
|  | R. Cerebellum (Lobule VIIa Crus II) | 26.3 | -81.9 | -35.3 | 197 | 3.69 | 0.27 |
|  | L. Angular Gyrus (TPJ) | -48.2 | -60.6 | 31.2 | 177 | 3.63 | 0.24 |
|  | Ventro-medial Prefrontal Cortex | -1.3 | 54.9 | -14.3 | 174 | 3.85 | 0.41 |
|  | L. Orbital Gyrus | -46.1 | 26.8 | -10.7 | 165 | 3.63 | 0.37 |
|  | Precuneus | 1.4 | -52.3 | 28.2 | 141 | 3.76 | 0.36 |
|  | R. Angular Gyrus (TPJ) | 62.7 | -47 | 23.4 | 127 | 3.64 | 0.22 |
|  | L. Cerebellum (Lobule VIIa Crus II) | -27.3 | -82.6 | -36.3 | 108 | 3.78 | 0.21 |
|  | R. Temporal Pole | 31 | 23.5 | -17.6 | 62 | 3.68 | 0.25 |
|  | L. Inferior Temporal Gyrus | -48.1 | 1.4 | -35.2 | 52 | 3.65 | 0.21 |
|  | R. Inferior Frontal Gyrus Pars Orbitalis | 52.8 | 28.6 | -4.7 | 50 | 3.57 | 0.32 |
|  | R. Angular Gyrus (TPJ) | 48.8 | -61.9 | 30.7 | 39 | 3.50 | 0.21 |
| Informative > Affective |  |  |  |  |  |  |  |
|  | R. Superior Parietal Gyrus | 12.8 | -67.9 | 51.5 | 99 | -3.79 | -0.17 |
|  | R. Angular Gyrus | 44.2 | -46.4 | 53.6 | 80 | -3.67 | -0.18 |
|  | R. Middle Frontal Gyrus | 44.5 | 42.2 | 17 | 58 | -3.65 | -0.21 |
| Informative > Direct |  |  |  |  |  |  |  |
|  | R. Superior Temporal Gyrus | 61.9 | -15.6 | -0.5 | 150 | 3.69 | 0.13 |
|  | R. Planum Temporale | 63 | -24.7 | 11.3 | 60 | 3.76 | 0.12 |

**FIG. 2.**
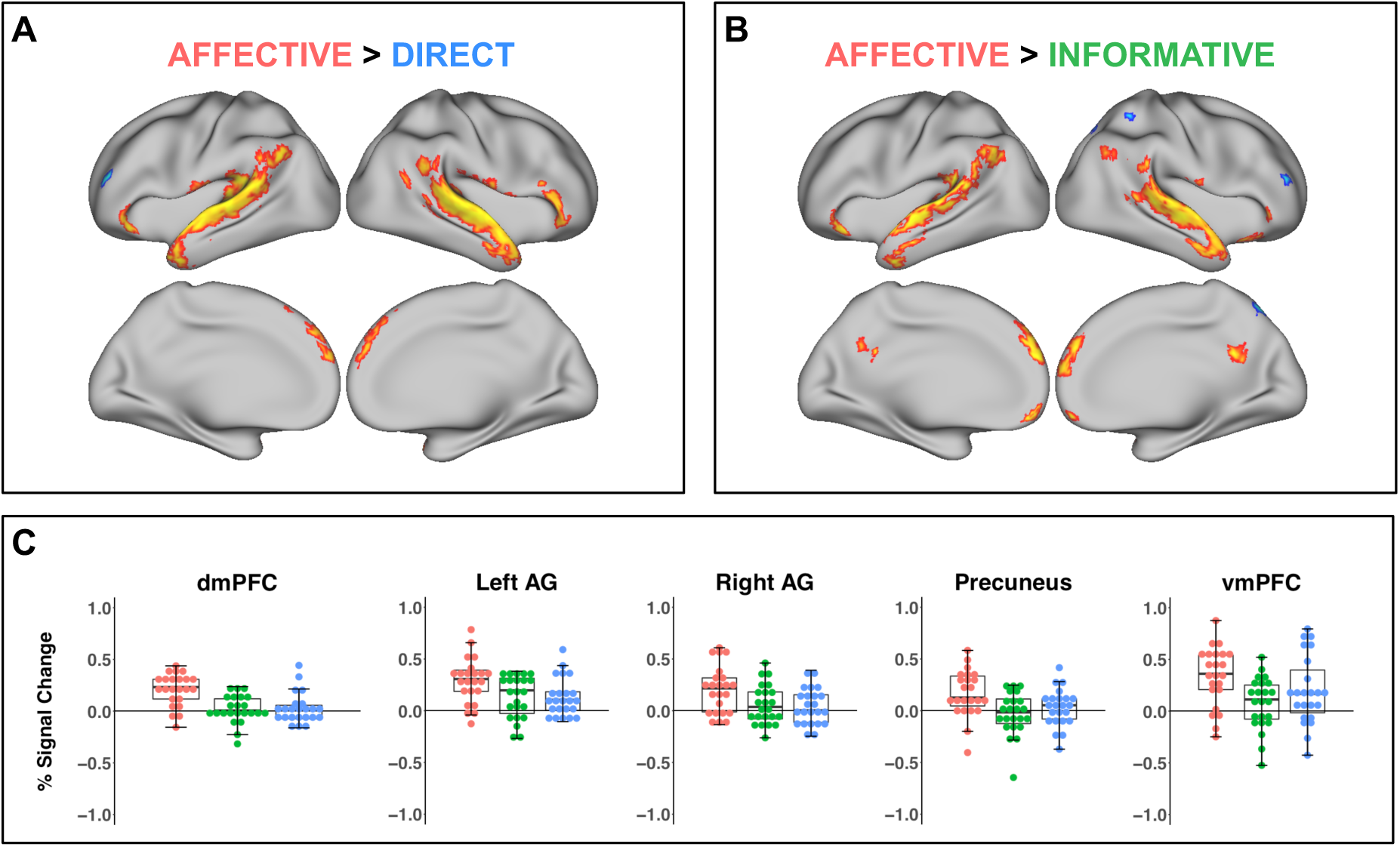
Significant clusters of activation for the (A) Affective > Direct and (B) Affective > Informative comparison displayed on an inflated cortical surface (voxelwise *p* < 0.001, clusterwise *p* < 0.05, FWE corrected). (C) Boxplots representing mean percent signal change per group and dot plots representing each participant in the affective (red dots), informative (green dots) and direct (blue dots) conditions for key regions identified in the comparisons between conditions: the dorsomedial prefrontal cortex (dmPFC), the left and right angular gyri (AG), the precuneus, and the ventromedial prefrontal cortex (vmPFC). Error bars represent standard error of the mean.

**FIG. 3.**
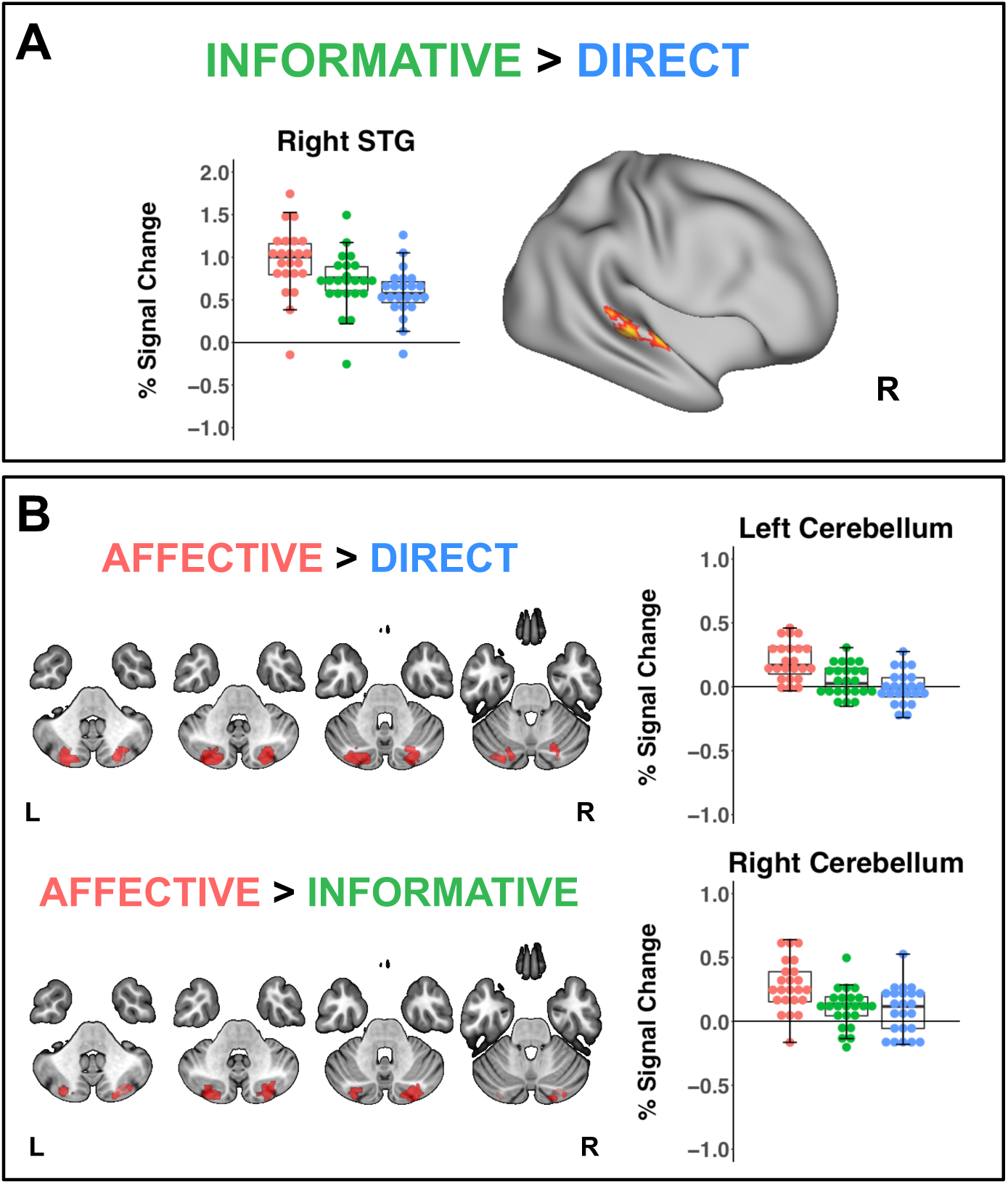
Significant clusters of activation for the (A) Informative > Direct comparison and (B) in the cerebellum for Affective > Direct and Affective > Informative comparisons (voxelwise *p* < 0.001, clusterwise *p* < 0.05, FWE corrected) with their respective boxplots and dot plots representing the mean percent signal change per condition. Error bars represent standard error of the mean.

## DISCUSSION

Our goal was to characterize the neural underpinnings of pragmatic language understanding in adolescent children. We chose to focus on indirect speech comprehension in discourse, specifically question/answer dialogues in which replies were the same across conditions but speaker meaning differed based on the context of the preceding question. Affective replies conveyed negative opinions, request refusals, or saving face (self or other’s); indirect informative replies added unsolicited information instead of a yes/no answer; and direct replies were literal answers to the question posed. The emotionally charged affective replies elicited the strongest response compared to informative and direct replies, primarily in ToM and language regions. This finding is consistent with the view of adolescence as a period of heightened sensitivity to social cues, especially peer acceptance and rejection cues [17]. Our study adds to the literature on the neural underpinnings of social cognition in adolescents and extends it to discourse and pragmatic language comprehension.

### Pragmatic language in the adolescent brain: the role of the language network

We hypothesized that indirect replies, informative and affective alike, would elicit higher activation in language processing areas than direct replies since both involve accessing the literal meaning and the intended meaning [4, 26, 47]. This hypothesis was supported by previous findings in adults showing bilateral STG activation for indirect vs. direct speech [5, 22]. While adolescent children showed increased bilateral activation in language regions when processing affective replies compared to direct replies, we did not find higher involvement in left temporal regions for informative replies. The only significant difference for informative vs. direct replies was found in the right STG/MTG and the right STG/PT, potentially reflecting speech and pitch/prosodic processing respectively. We speculate that the lack of more robust differences between the informative and the direct condition was due to: (1) shallow processing of informative replies, especially in the absence of an explicit pragmatic judgment task; (2) informative replies being as easy to process as direct replies. With respect to the second, evidence from corpora of natural speech show that indirect replies are usually given to questions requiring “polar” answers (yes or no) and that they are more frequent and more natural than direct replies-usually the interlocutor will anyway ask for more information than just a yes/no [61].

### Pragmatic language in the adolescent brain: the role of the Theory of Mind network

Adolescents showed higher activity in the mPFC, both angular gyri, and the precuneus, when processing affective replies as opposed to informative and direct replies. These areas are associated with perspective taking or ToM [69] and partly overlap with the default mode network (DMN) [35]. We propose that higher activity in these areas reflects perspective-taking processes [30], which are necessary to understand speaker intention in the affective dialogues. On the other hand, understanding speaker intention is not necessary when following informative or the direct dialogues, neither of which activated the ToM network, even when compared to baseline.

The comparison between affective and informative replies revealed two clusters in the mPFC: one dorsal and one ventral. Both regions are anatomically connected to the amygdala [28] and are associated with social cognition [8, 9]; the dmPFC is associated more with mentalizing and ToM while the vmPFC is associated more with reward, arousal, decision making [1, 18], and affective ToM [52]. Listening to dialogues in which one of the interlocutors is “politely” rejected or criticized could be salient enough to be perceived as arousing or threatening to adolescents for whom rewards and threats are primarily instantiated in the social sphere [17]. Adult listeners, on the other hand, consistently engage the dorsal part (dmPFC) for pragmatic language processing [5, 22, 27, 55, 56, 62]. Specifically, Basnakova et al. [6] found dmPFC activation for indirect (affective and informative) compared to direct replies and the anterior cingulate cortex for affective vs. informative indirect replies. Although we cannot make any developmental claims here, our findings are consistent with cross-sectional studies demonstrating that adolescents recruit anterior brain areas (such as medial prefrontal and inferior frontal) to a larger extent than adults when performing social reasoning tasks [11, 13, 52, 62]. Such a developmental pattern could be due to a combination of factors such as differences in brain maturation [58], consequences of hormonal changes during puberty (see [10] for a review), and/or differences in pragmatic competence.

We also found activation in the angular gyri bilaterally when processing affective compared to informative or direct replies. The clusters were localized primarily in cytoarchitectonic areas PGa and, partly, PGp of the angular gyrus, also referred to as posterior TPJ [36] or posterior angular part of the TPJ [42]. This area has been identified as part of a ToM network [35] while it is also considered part of the default mode network [46]. The TPJ is considered a hub for social cognition as it receives external, sensory information (dorsal part and STS part for face and biological motion), semantic information, and introspection/self-reflection and episodic memory information needed to construct situation models [42]. Unsurprisingly, given its overlap with social cognition, pragmatic language processing routinely engages the right [6], left [27, 47], or bilateral TPJ [5, 22, 56]. We observed that when EQ (which captured individual variation in social cognition abilities) or post scanning accuracy (which captured individual variation in attention and memory abilities) were added as covariates to the affective > informative and affective > direct replies comparison, only the left TPJ cluster remained significant. This result is in line with the suggestion by Amft et al. [1] that, while the TPJ in both hemispheres is involved in social cognitive processes, the left TPJ is additionally associated with language processing while the right TPJ with attention.

Informative indirect replies also elicited activity in the right AG when compared to affective indirect replies; however, this was a much more dorsal cluster, corresponding roughly to cytoarchitectonic area PFm (latIPS according to Patel et al. [42]) that extends to the superior parietal gyrus (BA 7) and the intraparietal sulcus. The connectivity profile of this part of the inferior parietal lobule is different from the more ventral TPJp [35], and it is associated more with information retrieval [54] and externally oriented attention [42] rather than social cognition. Lastly, we found higher activity in the precuneus for the affective > informative contrast. Although the precuneus has not been routinely considered part of a processing network for social cognition (see [8, 51]), its activation time course at rest correlates highly with that of the mPFC and the two angular gyri, forming the default mode network (DMN) [45]. Recent meta analyses have shown that the DMN overlaps with ToM areas in humans [1, 35] and macaques [35] and that the precuneus, together with the dmPFC, is associated with perspective taking, as well as self-referential and autobiographical processing [1]. When adolescents are tracking other people’s feelings, activity in the precuneus correlates with performance accuracy [30], and empathy training in adolescents results in strengthened connectivity between the precuneus and the vmPFC [31]. Finally, the precuneus is more engaged during indirect compared to direct speech processing in adult listeners [5, 22, 27, 55, 56].

### Activation beyond language and theory of mind related areas

We found differences in how affective and informative stimuli engage areas associated with attention and executive function. Affective stimuli elicited higher activity in the ventro-lateral prefrontal cortex (IFG) bilaterally, a finding consistent with findings in adults [6, 22] and most likely reflecting differences in working memory and inhibition processes [38]. The left IFG is part of a left fronto-parietal network whose time course tends to correlate with the DMN and introspective processes [32]. On the other hand, informative replies elicited higher activity in right fronto-parietal areas (primarily dorsal) compared to affective replies. Regions of the right fronto-parietal “task-positive” or dorsal attention network, which tend to be anti-correlated with DMN activity, are engaged during perceptual attention [32] and executive function processing [48] as well as reasoning [63]. Interestingly, higher activation in informative vs. affective replies in the right fronto-parietal areas was primarily driven by deactivation in the affective vs. baseline contrast - the informative vs. baseline contrast did not reveal significant change in these areas. In other words, the right fronto-parietal network was deactivated in the affective (most socially complex) condition while that was not the case for the informative condition.

Apart from cortical areas, affective replies elicited higher activity in the cerebellum bilaterally, specifically in lobule VIIa/Crus II. While often not imaged (not covered in many fields of view) or excluded from further analysis (see Feng et al. [22]), there is increasing evidence that the cerebellum is active during social cognition, working memory, semantic and pragmatic language tasks [24, 39, 47]. The cerebellum also shows increased connectivity with ToM/mentalizing network areas including the mPFC and the TPJ bilaterally during social cognition tasks [40].

While previous research has shown that adults engage the salience network when processing affective compared to informative replies, particularly in the anterior cingulate cortex and anterior insulae [6], this was not the case for our adolescent participants. The anterior cingulate cortex, anterior insula and premotor cortex, have been suggested to underlie “experience sharing” or affective empathy [69]. Given that, in the current study, experience sharing was neither necessary nor sufficient for understanding speaker intention, we did not anticipate activity in those areas. In fact, Kral et al. [30] found that when adolescents had to infer someone’s feelings, they were more accurate when they activated areas involved in perspective taking (including the mPFC, precuneus, superior temporal sulcus, and right temporoparietal junction) than areas relevant for experience sharing. The discrepancy between ours and previous findings might also be due to the fact that our dialogues did not include extensive contextual information which could have given rise to experience sharing as an epiphenomenon.

### Limitations and future directions

The main limitation of the present study is that it focuses on a single age group. Consequently, it cannot directly address developmental hypotheses but can only draw qualitative comparisons between current findings in adolescents and previous findings in adults with a similar paradigm [6]. Although similar, the two studies differ in two important ways: First, the studies differ in the language and culture of the participants, with one study performed on Dutch speakers living in the Netherlands, and the other on American English speakers living in the U.S. Different languages and cultures have qualitative differences in how indirectness or politeness is expressed as well as quantitative differences in how frequent indirect replies are [25, 65] (see also comments by Feng et al. [22] about natural indirect replies in Chinese). The second difference between the two studies relates to the design, most importantly the use of introductory context in Basnakova et al. [6] vs. no context in the current study. Given these discrepancies, future cross-sectional studies might help yield a better understanding of common and distinct neural underpinnings of indirect speech in children, adolescents and adults.

A second limitation concerns the differences between conditions in duration, speech rate and intonation. We tried to match the stimuli in the three conditions as closely as possible based on a variety of psycholinguistic measures. However, we opted for a naturalistic approach with respect to speech rate and intonation. As a result, affective dialogues differed from the informative and direct dialogues: questions and answers had more interesting intonation patterns and replies were slower and were preceded on average by a longer pause. Duration, speech rate and intonation were the few cues available to our listeners and, in the absence of any other context regarding the communicative situation, they undoubtedly relied on them to infer the intended speaker meaning.

Lastly, our participants were overhearers rather than addressees of indirect replies. In adult participants, being an overhearer or addressee of indirect face-saving replies does not elicit different responses in regions related to ToM [5]. However, we cannot exclude the possibility that, had they been the addressees rather than overhearers of indirect rejections or refusals, adolescent listeners would have shown different activation patterns, potentially engaging limbic and salience network areas. Future work is needed in order to empirically test this possibility.

## CONCLUSIONS

This is, to the best of our knowledge, the first study to look at the neural basis of pragmatic language processing in adolescent children. We demonstrated that adolescents show more widespread brain activity, particularly in regions related to theory of mind and language, when processing indirect speech that conveyed negative opinions, refusals or face-saving, than when it simply added more information in response to a question. We interpret these results in the light of accumulating evidence that adolescents show heightened sensitivity to social and affective information compared to adults, extending this body of evidence to include pragmatic language.

## ACKNOWLEDGMENTS

We thank Kristi Schonwald, Hannah Barton, Marisha Kazi, Kelly Walbert, Jessica Odbert, Todd Parish, Azmi Banibaker, Inez Falcon-Haus, Danny Siu, and the children and families who participated in the study. We are also grateful to Bruno Bara, Jana Basnakova, and Rósa Gísladóttir. This research was supported by the National Institutes of Child Health and Human Development (Grant P01 HD 40605).

